# Efficient calculation of orientation-dependent lipid dynamics from membrane simulations

**DOI:** 10.1101/2023.05.23.542012

**Authors:** Milka Doktorova, George Khelashvili, Michael F. Brown

## Abstract

Molecular dynamics simulations of lipid membranes have become increasingly impactful in biophysics because they offer atomistic resolution of structural fluctuations in relation to their functional outputs. Yet quantitative characterization of multiscale processes is a formidable challenge due to the distribution of motions that evade analysis of discrete simulation data. Here we investigate the efficient calculation of CH bond relaxation rates from membrane simulations. Widely used computational approaches offer numerical simplicity but fall short of capturing crucial aspects of the orientation dependence of the dynamics. To circumvent this problem, we introduced a robust framework based on liquid crystal theory which considers explicitly the CH bond motions with respect to the director axis (bilayer normal). Analysis of the orientation dependence of the dynamics shows excellent agreement with experiment, illustrating how the ordering potential affects the calculated relaxation rates. Furthermore, a fit-based resampling of the autocorrelation function of the bond fluctuations validates the new approach for low-temporal resolution data. The recovered relaxation rates indicate that at short timescales, both with and without cholesterol, the local motions of CH bonds describe the bilayer microviscosity and resemble liquid hydrocarbons. Our results establish the critical role of the orientational anisotropy in analysis of membrane simulations, explain fundamental aspects of lipid dynamics, and provide guidelines for extracting information that can be compared to experimental data.

**STATEMENT OF SIGNIFICANCE:** Nuclear magnetic resonance data have been historically used to validate membrane simulations through the average order parameters of the lipid chains. However, the bond dynamics that give rise to this equilibrium bilayer structure have rarely been compared between in vitro and in silico systems despite the availability of substantial experimental data. Here we investigate the logarithmic timescales sampled by the lipid chain motions and confirm a recently developed computational protocol that creates a dynamics-based bridge between simulations and NMR spectroscopy. Our results establish the foundations for validating a relatively unexplored dimension of bilayer behavior and thus have far-reaching applications in membrane biophysics.

## INTRODUCTION

Functions of membrane-associated molecules are often inextricably coupled to the structural and dynamical properties of the lipid bilayer matrix. Various experimental techniques have been transformative for probing the conformational and energetic landscapes of membrane lipids and their dependence on composition, temperature, and pressure variables (1-7). Notably small-angle scattering (SAXS and SANS) data characterize bilayer thickness and area packing per lipid (8); fluorescence microscopy (9), resonance energy transfer (FRET) (10), and cryogenic electron microscopy (cryo-EM) (11,12) reveal phase coexistence in lipid mixtures; flicker spectroscopy (13,14) and neutron spin-echo measurements (15-18) report on bilayer elasticity; and fluorescence correlation spectroscopy quantifies lipid diffusion in various biomedical contexts (19,20). While experimental results have shaped our comprehension of lipid properties over a wide range of length- and time scales, the inability to access individual molecules has created room for computational and theoretical investigations (21-27). In particular, molecular dynamics (MD) simulations have uncovered otherwise inaccessible mechanisms occurring at the nanoscale (21-23,28,29), including the role of membrane deformation in peptide-mediated lipid flip-flop between leaflets (30), the effect of cholesterol on electrostatics-driven protein binding to membranes (31), contributions of interleaflet coupling to phase separation (32), and the pathway of spontaneous lipid translocation between opposite leaflets (33). Often perceived as a computational microscope, these MD studies have enabled the unifying comparison of different experimental techniques. Moreover, they have guided the refinement of data-driven models critical for analysis of experiments using the next generation of physics-based force fields (8,11,34,35).

Clearly the versatility and applicability of molecular simulations hinges upon the robust validation of the trajectories against increasingly stringent experimental data (2,15,36-38). Studies related to membrane structure and dynamics rely heavily on the accurate parameterization of the interatomic and intermolecular interactions (28,39-41). Such sets of parameters or lipid force fields, developed for both all-atom and coarse-grained molecular models, govern the bilayer equilibrium and dynamical properties and have been optimized against bilayer structural parameters obtained mostly from scattering (SAXS and SANS) and solid-state NMR measurements (42,43). Ongoing refinements of these lipid force-field parameters have proven successful in reproducing certain aspects of the experimental data, while missing others, affirming the inherent need for more points of comparison (34). In that respect, investigations of membrane dynamics, in addition to average structure, give a promising yet relatively less explored fourth dimension of lipid biophysics. Recently we showed how the NMR relaxation rates of lipids in lamellar samples can be directly compared to the carbon–hydrogen (CH) bond motions in simulated bilayers through mapping the spectral densities of the fluctuations (23). Analogous calculations have been done in the past using the widely applied status quo approach developed for isotropic motion in nonlamellar systems. Here we show that this classical methodology while useful misses a crucial feature of CH bond motions—namely, the angular anisotropy relative to an external frame. As a result, it provides only approximate results for the bond relaxation rates.

In the present article, we extend our approach to quantify the angular dependence of lipid dynamics from the simulation trajectories, obtaining results in excellent agreement with experimental measurements. This dependence of lipid fluctuations on the bilayer orientation is explicitly incorporated into our recently developed computational protocol (23). While quantifying the CH bond reorientations demands only knowledge of changes in atomic positions over time, calculating the respective spectral densities entails a Fourier transform of the autocorrelation function of the fluctuations. This calculation turns out to be particularly challenging due to the discrete nature of the multiscale simulation data. We describe the problem and its origins (35,44-46), and demonstrate the validity of a solution that circumvents the sampling issues for data with relatively low temporal resolution (23). For the current application, the simulated relaxation rates are explained by collective lipid motions and membrane elasticity, while the bilayer core resembles liquid hydrocarbons in agreement with experiments.

## THEORETICAL METHODS

### Simulation protocol

We investigated the properties of fully atomistic lipid bilayers: 1,2-dimyristoyl-*sn*-glycero-3-phosphocholine (DMPC) containing 0, 33, and 50 mol% cholesterol. Each bilayer contained 100 lipids per leaflet (200 lipids total) and 45 water molecules per lipid with no added salt ions. The bilayers were simulated in the NPT ensemble at a constant temperature of 317 K (44°C). The original 2-μs-long trajectories, performed with OpenMM (47) and the CHARMM36 force field for lipids (28), were taken from (23). These simulations employed a timestep of 2 fs and atomic coordinates were saved every 40 ps. To analyze the dynamics at shorter length scales, we resampled the DMPC trajectories with 0 and 33 mol% cholesterol by restarting the simulation at different time points and running short simulations with more frequent data output. In particular, starting at 0.8, 1.0, 1.2, 1.4, 1.6, 1.8, and 2.0 μs of the original trajectories we ran 400-ps-long simulations by using stored restart files of both atom positions and velocities, and outputting atomic coordinates every 10 fs. Except for the output frequency, these resampling simulations were performed with the same simulation parameters as the original long trajectories as described in detail in (23) including a timestep of 2 fs with *constraints* set to HBonds, a 10–12 Å potential for van der Waals interactions, and Langevin dynamics maintaining a temperature of 317 K with a friction parameter of 1.0 per picosecond. In addition, starting from the ends of all three 2-μs-long trajectories, we ran additional 800-ns-long simulations, again with the same simulation parameters but with an output interval of 4 ps. The data were used to evaluate the accuracy of the mathematical resampling performed on data with lower temporal resolution as a way to recover the unbiased relaxation rates.

### Representation of carbon-hydrogen bond orientations in Cartesian coordinates

The fluctuations of the carbon-hydrogen (CH) bonds of the lipid acyl chains were quantified as in (23). Briefly, the instantaneous orientation of a CH bond is described by the Euler angles *Ω* = (*α, β, γ*) = (*α*_PD_, *β*_PD_, *γ*_PD_) in the director frame (Fig. 1A) where *α* = 0 due to axial symmetry about the CH bond axis, β defines the angle that the CH bond (<u>P</u>rincipal axis) makes with the bilayer normal (<u>D</u>irector axis, **N**_B_) along the *z*-dimension of the simulation box, and *γ* quantifies the CH bond rotation around the director. The orientation of a CH bond at time *t* is then described in 3-dimensional space by functions of *β* and *γ*, the so-called Wigner rotation matrix elements (48), given by:

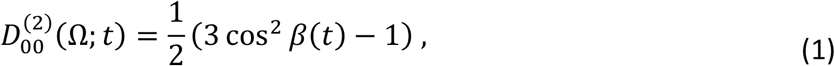

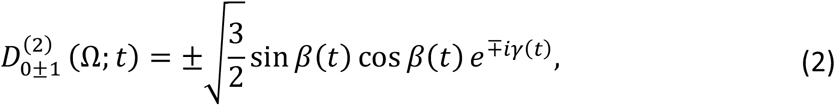

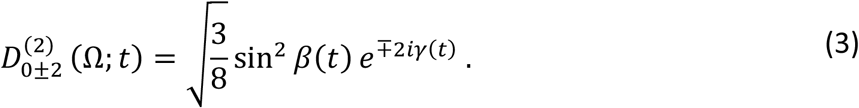

where 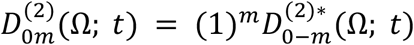 with *m* being 0, ±1, or ±2, and the right-hand side indicating the complex conjugate. The above functions are evaluated for every CH bond in the bilayer and averaged over the respective carbon atoms across all lipids.

**Figure 1.**
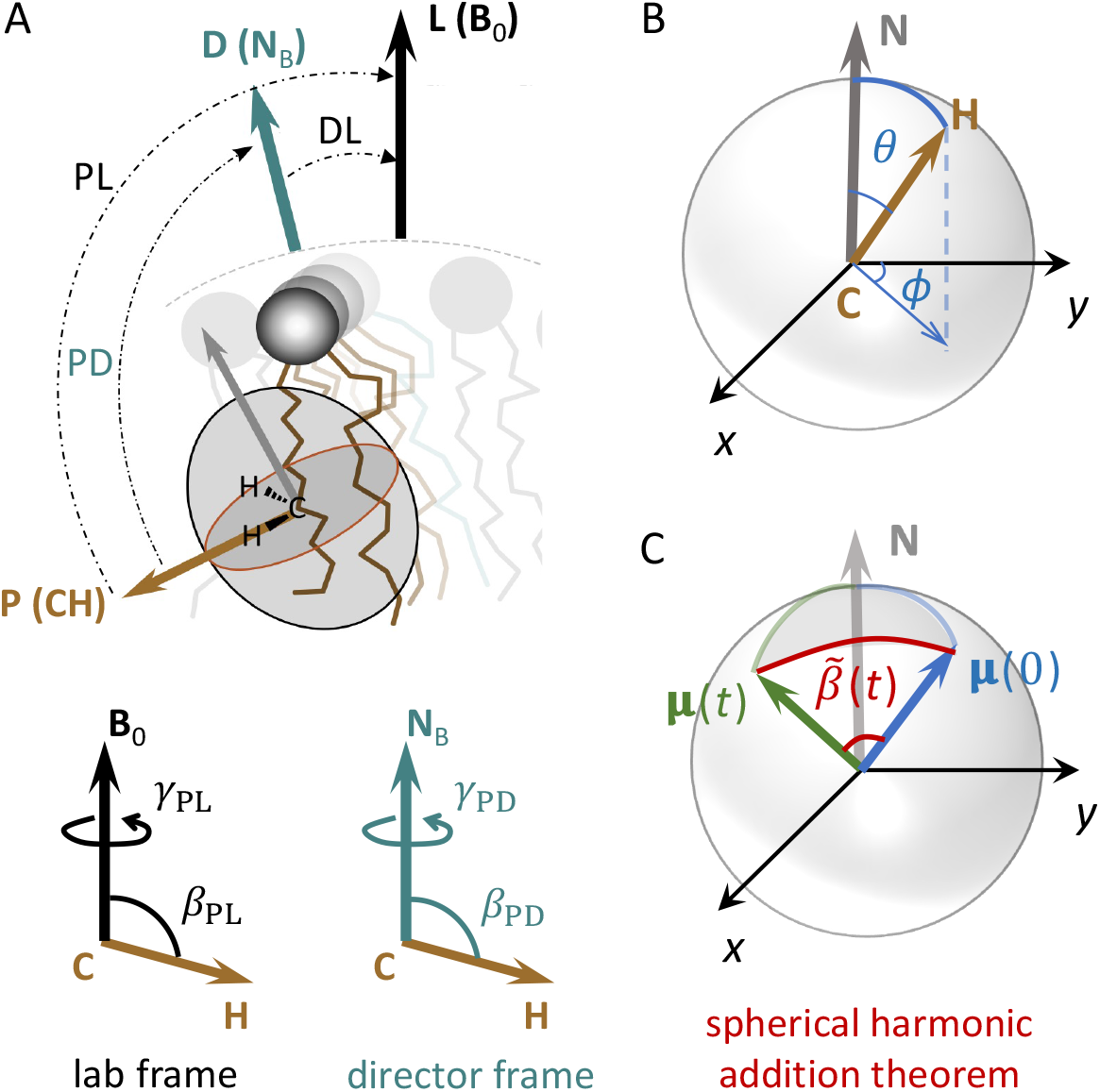
Schematic illustration of angles and frames of reference used to analyze lipid dynamics. (A) Representation of a carbon–hydrogen bond on a lipid chain and the Euler angles *β* and *γ* in different frames of reference. These angles can describe orientations in the laboratory (lab) frame (*β*_PL_, *γ*_PL_), i.e., between the principal (CH bond) axis and the fixed magnetic field or laboratory (**B**_0_) axis; or in the director frame (*β*_PD_, *γ*_PD_) between the principal (CH bond) axis and the director (bilayer normal, **N**_B_) axis. The *β* angle is the angle that the CH bond makes with the main-frame axis, and the angle *γ* defines the rotation of the CH bond around that axis which is calculated as described in (23). (B) The CH bond orientation with respect to any axis **N** can be defined in an analogous way in spherical polar coordinates with the polar angle *θ* and azimuthal angle *ϕ*. (C) Using the spherical harmonic addition theorem, the time-dependent reorientation of the CH bond is described with its positions at time 0, given by **μ**(0), and time *t*, given by **μ**(*t*). The angle between these two vectors is 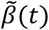 which is different from the angles *β*_*PD*_ and *β*_*PL*_ in (A) [or *θ* in (B)] as it does not depend on the direction of the **N** axis. **[1-column figure]**

### Mean director-frame relaxation rates from Wigner *D*-functions

As described in (23), the relaxation rate *R*_1Z_ of the CH bond fluctuations is obtained by first calculating the correlation function of the fluctuations, then performing a Fourier transform to obtain the spectral density, and evaluating it at ω_0_ and 2ω_0_ where ω_0_ is the Larmor frequency of the NMR measurement. Since the fluctuations are described by separate Wigner rotation matrix elements (Eqs. 1–3) and each of them has its own correlation function, *R*_1Z_ is a linear combination of the corresponding spectral densities at the frequencies of interest. In simulations, the measured fluctuations (and relaxation) are with respect to the bilayer normal **N**_B_, that is, the director frame, while in NMR experiments, they are always with respect to the fixed magnetic field axis **B**_0_, i.e., in the laboratory frame (Fig. 1A). This means that if **N**_B_ is parallel to **B**_0_, the results from the two techniques would have perfect correspondence. However, experiments are often done on liposome samples in which **N**_B_ adopts all possible orientations relative to **B**_0_ due to the spherical geometry of the vesicles. Therefore, connecting the results from the simulations to the actual experimental data requires transforming or averaging of the latter. Below we outline the steps of this transformation, which is described in more detail in the supplemental material (SM) of (23).

We will refer to the coordinate system (*X, Y, Z*) as the laboratory (lab) and start by expressing the CH bond fluctuations directly with respect to the lab frame. In that case, the correlation function 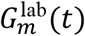, spectral density 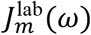, and relaxation rate *R*_1Z_ can be all written as:

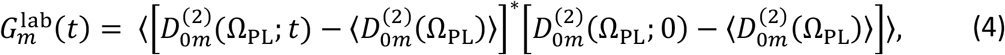

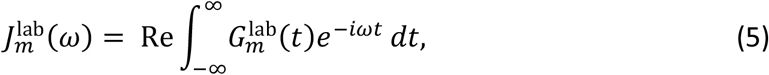

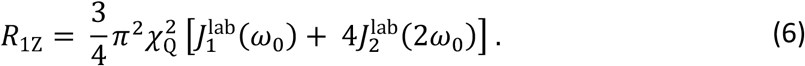

In Eqs. 4–6, Ω_PL_ = (α_PL_, *β*_PL_, *γ*_PL_) denotes the Euler angles between the CH bond (<u>P</u>rincipal axis) and the magnetic field **B**_0_ (<u>L</u>aboratory axis), see Fig. 1A. The subscript *m* is 0, ±1, or ±2, *t* is time, and ω is frequency with ω_0_ being the Larmor frequency of the measurement. In Eq. 4 the 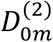 matrix elements are as defined in Eqs. 1–3 and the angular brackets ⟨…⟩ denotes a time or ensemble average. In Eq. 6, the static quadrupolar coupling constant, χ_Q_ ≡ *e*^2^*qQ/h* = 170 kHz, and the value of the numerical pre-factor is (3/4)π^2^(1.70 × 10^5^)^2^ = 2.1<u>39</u>2 × 10^11^ *s*^2)^. The reader should recall that for isotropic liquids there is no projection index, so that *J*_*m*_(ω) → ⟨*J*_*m*_(ω)⟩ ≡ *J*(ω) = (1/5)*j*(ω) where *j*(ω) is the reduced spectral density (see Eq. S20 in the SM of (23)).

In a simulation, the coordinate system is defined by the bilayer director **N**_B_, i.e., (*x, y, z*) and we will refer to it as the director frame. Here, the correlation function 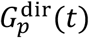 and spectral density 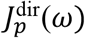 can be analogously written as:

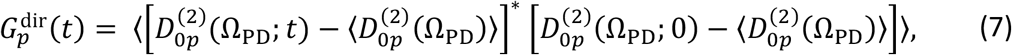

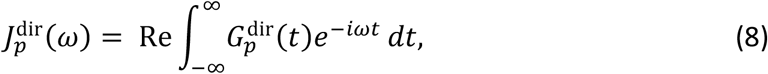

where Ω_PD_ = (*α*_PD_, *β*_PD_, *γ*_PD_) denotes the Euler angles with respect to the director frame (Fig. 1A). Note that here again the subscript *p* is a projection index that can be equal to 0, ±1, or ±2. It follows that Eqs. 7 and 8 can be calculated from simulations and to connect with experiment, all we need is to write *R*_1Z_ from Eq. 6 as a function of 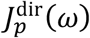 using Eq. 8. This can be achieved by following the principle of closure, where if Ω_DL_ = (*α*_DL_, *β*_DL_, *γ*_DL_) denotes the Euler angles between the <u>d</u>irector axis **N**_B_ and the laboratory frame axis **B**_0_ (Fig. 1A), then:

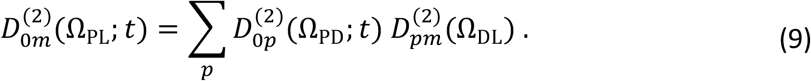

Substituting Eq. 9 into Eq. 4 leads to

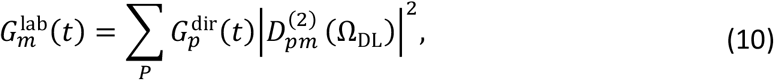

and likewise, inserting Eq. 10 into Eq. 5 yields:

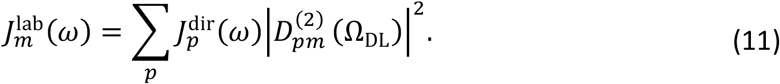

Note that in Eqs. 10 and 11 it is assumed the motions of the lipids are cylindrically (rotationally) symmetric about the director axis.

Since in a typical liposome sample **N**_B_ adopts all possible orientations relative to **B**_0_ on the timescale of the NMR measurement, the mean-square Wigner rotation matrix elements for the transformation from the director frame to the lab frame are averaged to their isotropic values, leading to 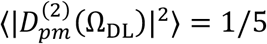. The factor of 1/5 comes from considering the spherical (“powder”) averaging as part of a 2-step process: first, the CH bond rotation with respect to the director axis, and then rotation about the director axis versus the magnetic field axis. This makes the right-hand side of Eq. 11 independent of *m*, corresponding to the mean director-frame spectral density. Because the summation is over all *p* ∈ [0, ±1, ±2] and 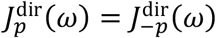, we can explicitly write that:

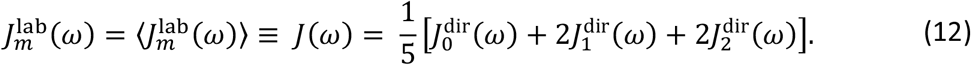

Lastly, substituting Eq. 12 into Eq. 6 gives us an expression for the orientationally averaged experimental *R*_1Z_ rate as a function of the computationally accessible 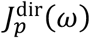 spectral densities:

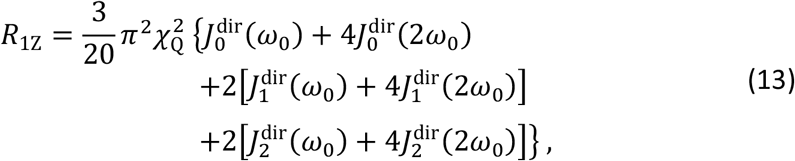

where the pre-factor is equal to (3π^2)^⁄20)(1.70 × 10^5^ s^−1^)^2^ = 4.2<u>7</u>85 × 10^10^ s^−2)^.

### Orientation-independent relaxation rates from spherical harmonics

In addition to expressing *R*_1Z_ as a function of the director-frame spectral densities 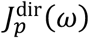 as in Eq. 13, the experimentally measured value from Eq. 6 can be approximated by an orientation-independent relaxation rate. As mentioned above, due to the distribution of the bilayer normal in a liposome sample, the spectral density 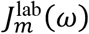 in the laboratory frame can be orientationally averaged to its isotropic limit *J*(ω), which does not depend on the projection index *m* (Eq. 12). Analogously, the correlation function 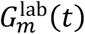 from Eq. 10 can be written as:

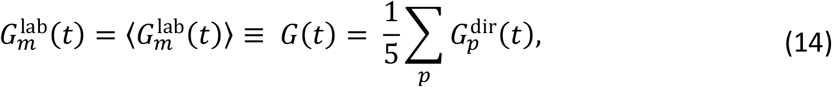

where the angular brackets denote a time or ensemble average. In Eq. 14 *G*(*t*) represents the mean director-frame correlation function, which corresponds to an anisotropic liquid-crystal model. The factor of 1/5 comes from the orientational average of the director (bilayer normal) as mentioned above.

As described in the supporting material (SM) of (23), the correlation functions 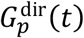, which in Eq. 7 are functions of the Euler angles in Cartesian coordinates, can be expressed in terms of spherical harmonics, 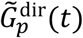, as follows:

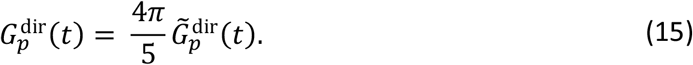

In Eq. 15 and throughout, the tilde refers to functions of the spherical harmonics that are distinguished from those using Wigner *D*-functions. From here, the mean director-frame correlation function *G*(*t*) from Eq. 14 becomes:

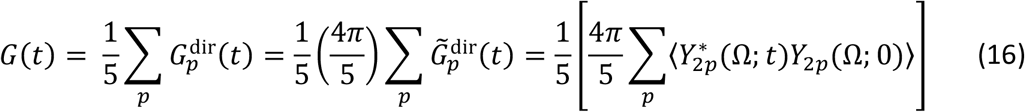

where the *Y*_2*p*_(Ω) functions are the spherical harmonic equivalents to the 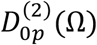 functions:

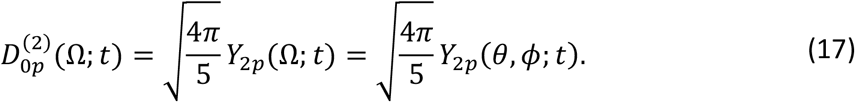

Note that in Eq. 17 the orientation of the CH bond can be represented using either the Euler (Ω; Fig. 1A) or spherical polar (*θ, ϕ*; Fig. 1B) angles as the basis, with the correspondence shown in Fig. S1 of (23).

Next, if we now assume spherical symmetry with no dependence on the specific orientation of the CH bond with respect to **N**_6_ (or **B**_0_), then we can apply the spherical harmonic addition theorem to simplify the expression in the square brackets in Eq. 16, yielding:

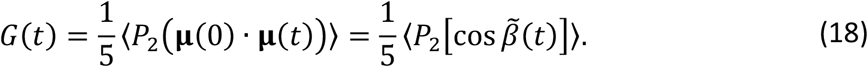

The reader should take note that in Eq. 18, the second-order Legendre polynomial *P*_2_ has the same functional expression as 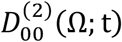 in Eq. 1, where 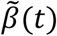 is the angle between the CH bond at time 0 and time *t*. In other words, if the unit vector *μ*(*t*) describes the direction of the CH bond at time *t* as shown in Fig. 1C, then the reorientation of the bond in Eq. 18 is defined by the scalar product of *μ*(0) and *μ*(*t*), i.e., the cosine of the angle 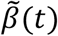 between the two vectors. Because the scalar product is invariant, the spherical harmonic addition theorem eliminates the dependence of the correlation function on the direction of the bilayer normal **N**_B_ (or laboratory axis **B**_0_) by considering the *change* in *β* described by 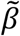 (Fig. 1C), and not *β* itself (Fig. 1A). Whether this is actually the case for lipid membranes is further discussed below.

Notably in Eq. 18, the mean *P*_2_ function of cos 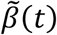 is typically denoted as *C*(*t*) (49), where the corresponding spectral density *J*_K_(ω) is written as:

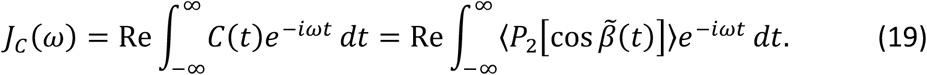

The above formula corresponds to a 1-step motional process, i.e., the CH bond motion is considered with respect to an arbitrary axis. From Eq. 19 we then obtain the orientation-independent relaxation rate in terms of the computationally accessible spherical harmonics, which reads:

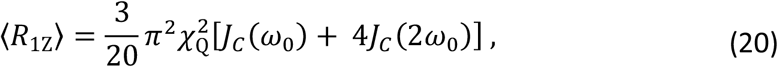

where *J*_K_(*ω*) is the two-sided Fourier transform of the *C*(*t*) correlation function. The above expression, Eq. (20), is the same as in the textbook case for solution NMR spectroscopy (50) with the substitution *J*(*ω*) → (1/5)*J*_K_(*ω*) (compare Eqs. 6 and 20 above). Because the director axis is not included, the orientational (“powder”) average is attributed entirely to the CH bond motion, e.g., as in (but not restricted to) the Debye model for rotational relaxation of an isotropic liquid. Inclusion of an alignment frame (e.g., director) leads to the generalized model-free (GMF) approach as originally described (44).

### Calculation of experimental relaxation rates from molecular simulations

Next, we can calculate 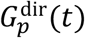 and 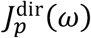 from Eqs. 7 and 8, using the simulation trajectories as described in (23). Briefly, for each of the time series, the element 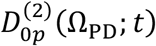 can be written as 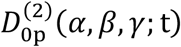 where *p* ∈ [0,1,2], the angle *α* = 0, and we drop the subscript PD for convenience, since all Euler angles in flat bilayer simulations are with respect to the director frame whose *z*-axis is the membrane normal (Fig. 1A). For every carbon 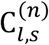 where 1 ≤ *l* ≤ *N*: is an individual DMPC lipid (*N*_L_ being the total number of DMPC lipids in the bilayer), and 2 ≤ *n* ≤ 14 is the carbon number on the *sn*-1 (*s* = 1) or *sn*-2 (*s* = 2) chain, the autocorrelation function can thus be obtained from:

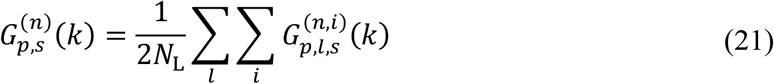

with

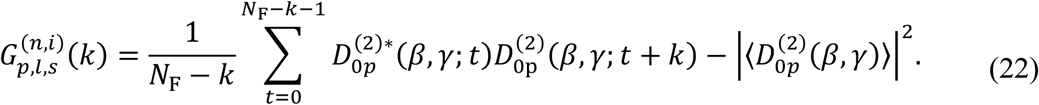

In Eq. 21 the inner summation is over the two hydrogen atoms *i* = (1,2) at carbon 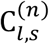 and in Eq. 22 the second term is the squared mean of the fluctuations; *k* is the lag time, *N*_F_ is the total number of trajectory frames (or time points), and 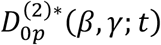 denotes the complex conjugate of the 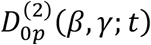 rotation matrix element. At zero lag time, i.e., *k* = 0, the autocorrelation function in Eq. 22 yields the variance of 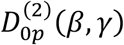, which reads:

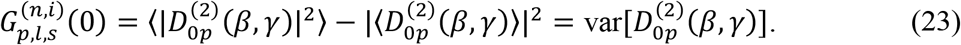

From Eq. 22 the spectral density function of the fluctuations is the two-sided Fourier transform (FT) of the autocorrelation function. Numerically however we can calculate only the single-sided FT, so that we can estimate the double-sided FT as follows:

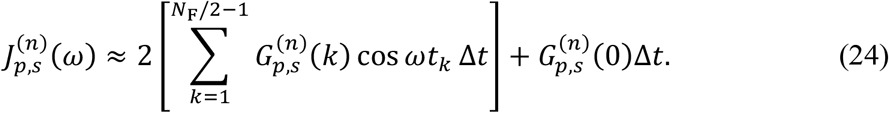

Note that in Eq. 24, the frequency *ω* is related to the Larmor frequency *v*_0_ of the NMR instrument by *ω*_0_ = 2π*v*_0_, *Δt* is the sampling time interval, and *t*_*k*_ = *kΔt* is the time at lag *k*. The discrete spectral density is fit to either a power-law function (if a fit of the correlation function is used, see Results) or to a simple smoothing spline function to access values at specific frequencies. Following Eq. 13, the relaxation rate is then calculated as:

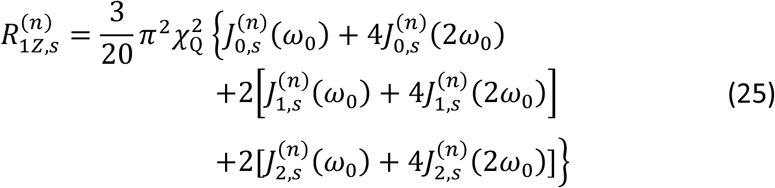

with the pre-factor equal to (3π^2^⁄20)(1.70 × 10^5^ s^−1^)^2^ = 4.2<u>7</u>85 × 10^10^ s^−2^. The relaxation rate in Eq. (25) represents an average over all director orientations as discussed above, and we refer to it as the *mean director-frame relaxation rate*.

To calculate the orientation-independent relaxation rate with Eq. 20, we first need to compute the two-sided Fourier transform of the *C*(*t*) correlation function. Traditionally, this is done by using the single-sided FT, *j*_K_(*ω*) and multiplying the result by 2 which gives:

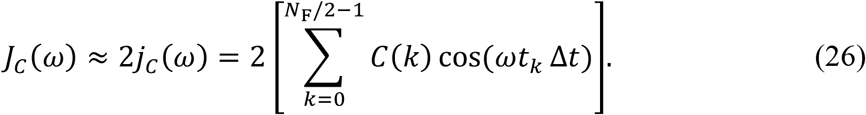

The corresponding orientation-independent relaxation rate is then written as:

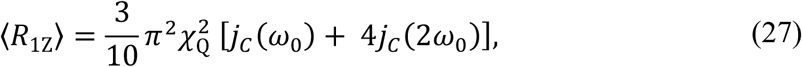

in which the pre-factor is equal to 8.5<u>5</u>7 × 10^10^ s^−2)^ and is the same as the pre-factor in Eq. 2.5 in (51).

Analogously to Eq. 24, the spectral density *J*_K_(*ω*) can be approximated from the single-sided FT by ensuring that the zeroth element of the correlation function, *C*(0) = 1, is not counted twice. We will refer to the resulting relaxation rate as the mean *corrected* orientation-independent relaxation rate ⟨*R*_1Z_⟩_corr_:

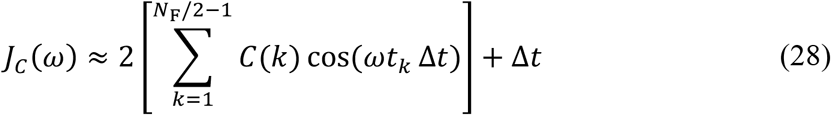

in which

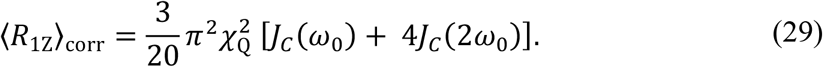

Here the pre-factor in Eq. 29 is equal to 4.2<u>7</u>85 × 10^10^ s^−2^. In both cases (Eqs. 26 and 28), the spectral density is fit to a smoothing spline function to estimate the values at *ω*_0_ and 2*ω*_0_.

### Calculation of effective correlation times

In our approach, the correlation times of the CH bond fluctuations are described by the functions in Eqs. 1–3 and can be obtained from the corresponding autocorrelation functions 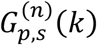 with Eq. 21. However, in general a broad distribution of correlation times is to be expected for either collective or noncollective lipid bilayer motions (44). For a more recent discussion please see Refs. (35,52). In the case of lipid motions, we follow the procedure of Giovanni Lipari and Attila Szabo (49) based on Padé approximants (53), where the effective correlation time, τ_eff_, is related to the integral of 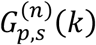 as follows:

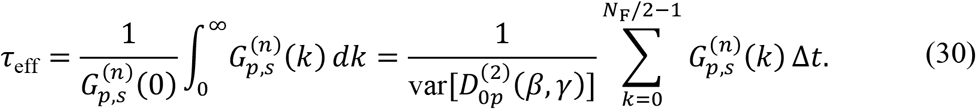

### Analysis of isomerization rates

For all consecutive 4-carbon segments along the lipid chains, the isomerization rates were analyzed following the approach outlined in (54). First, the dihedral angles of all segments were calculated and used to determine the cutoff angle that classifies the segment conformation as *trans* versus *gauche* (±120 degrees). Then for each segment a time series *N*_A_(*t*) was constructed such that in every frame (i.e., at every time point *t*) a value of 1 or 0 was assigned to *N*_A_(*t*) depending on whether the segment had a *trans* (0) or *gauche* (1) isomerization. The resulting number correlation function *C*_N_(*t*) was calculated from δ*N*_A_(*t*) = *N*_A_(*t*) − ⟨*N*_A_ ⟩ following Eq. 7a from (54) with the MATLAB *xcorr* function specifying the “normalized” option which ensures that *C*_N_(0) = 1. The correlation functions for all lipids and in all frames were averaged, and Eq. 30 was used to calculate the effective correlation time for the segment. Note that due to the noise the averaged correlation functions decayed to a value slightly different from 0, so that the mean of the last quarter of the correlation function was subtracted from *C*_N_(*t*) prior to calculation of the τ_eff_ correlation time. This quantification of the carbon segment dynamics makes them directly comparable to the effective correlation times of lipid CH bond fluctuations.

## RESULTS

### Molecular simulations capture the angular dependence of bond relaxations and yield orientationally averaged relaxation rates

Knowing what exactly gives rise to the relaxation rates measured in a solid-state NMR (ssNMR) experiment is essential for properly translating data to MD simulations. In the context of CH bond fluctuations, there are two main frames of reference in the experiment: the laboratory (or lab) frame defined by the fixed magnetic field axis **B**_0_, and the director frame specified by the normal to the bilayer surface, **N**_B_ (Fig. 1A). Any quantity measured with ssNMR is in the lab frame, that is the dynamics are always with respect to **B**_0_. The CH bonds of lipids, for example, have restricted fluctuations in the director frame, and the ordering potential induced by the **N**_B_ director gives rise to the well-known order parameters of the fluctuations (1,55). The order parameter is an average property measured from the NMR lineshape that comes from the reorientation of all lipid CH bonds relative to the lab frame **B**_0_, e.g., as described by a mean-torque model (56).

For a typical liposome sample, the bilayer directors adopt all orientations with respect to the fixed axis of the magnetic field. This orientation of the directors does not affect the average order parameters; however, it is relevant for interpretation of the *R*_1Z_ relaxation rates of the CH bonds. That is due to the fact that the experimentally measured *R*_1Z_ value depends on the angle that the bilayer director makes with the magnetic field axis. This was shown in work with oriented bilayer samples in the 1990s by various research groups (57-60). There the bilayers were gradually rotated with respect to **B**_0_ yielding distinct changes in the measured relaxation rates. This angular dependence is not apparent in relaxation measurements of multilamellar lipid dispersions but is clearly visible in data from microcrystalline powders (61). Specifically, Brown and Davis (62) were able to show experimentally that the relaxation anisotropy present in multilamellar vesicles (MLVs) is orientationally averaged by lipid translational diffusion, which happens on timescales shorter than those of the actual spectral measurements.

The orientational distribution of the directors in liposome samples can be modeled as uniform on the surface of a unit sphere due to the geometry of the vesicles. Here we consider a solid angle Ω defined by spherical polar coordinate (*θ, ϕ*) with the differential solid angle *d*Ω = sin*θdθdϕ*, where *θ* is the polar angle (colatitude or zenith) and *ϕ* is the azimuthal angle (longitude) in the *x-y* plane from the *x*-axis. Conservation of probability then leads to *P*(Ω)*d*Ω = *P*(*θ, ϕ*)*dθdϕ* for the infinitesimal probability, in which the probability density function is given by *P*(*θ, ϕ*) = sin *θ*/4π. For a fluid lipid bilayer with axial symmetry about the bilayer normal (director), the azimuthal probability density entails integrating *P*(*θ, ϕ*) over the polar angle *θ* in the interval *θ* ∈ [0, π] yielding *P*(*ϕ*) = 1/2π. Alternatively, we can integrate *P*(*θ, ϕ*) over the azimuthal angle *ϕ* in the interval *ϕ* ∈ [0,2π) to get the probability density for the polar angle. The normalized probability density function is thus mathematically described by

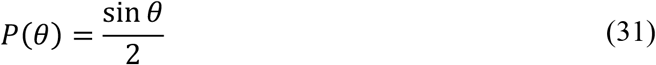

in which *θ* ∈ [0, π] is the polar angle, with *d*Ω = sin*θdθdϕ* for the differential solid angle. As illustrated by Eq. 31, the uniform distribution on a unit sphere does not imply equal probability but instead, orientations with *θ*∼90° are most likely. The measured relaxation rate is then an orientational average of contributions from the various directors, that is, the sum of relaxation rates for the distinct orientations, each scaled by the probability *P*(*θ, ϕ*) of that orientation. Take note that *P*(*θ, ϕ*) is invariant with respect to the *ϕ* angle due to the spherical symmetry.

One way to directly relate the experimental results to numerical simulations is to first estimate the relaxation rate from the MD trajectories in the laboratory frame. Simulated bilayers are usually flat membrane patches with a single bilayer director parallel to the *z*-dimension of the simulation box. In that respect, they resemble oriented bilayers that can be rotated to examine the angular dependence of the relaxation. Accordingly, we analyzed two fluid bilayer trajectories from (23), DMPC with and without 50% cholesterol, simulated at 44°C. We assumed that **B**_0_ is initially parallel to the bilayer normal **N**_B_, i.e., the angle *θ* between **B**_0_ and the *z*-dimension of the simulation box was 0 degrees. The **B**_0_ axis was then gradually rotated by increasing *θ* as shown in Fig. 2A, and the relaxation rates calculated directly with respect to **B**_0_, i.e., in the lab frame using Eq. 6. Rotating **B**_0_ relative to **N**_B_ is analogous to keeping the orientation of **B**_0_ fixed and rotating the bilayer patch, as done in the actual NMR experiment. Indeed, using this approach we were able to recover almost perfectly the angular dependence observed experimentally with solid-state NMR spectroscopy (Fig. 2B–C, red and black symbols). This dependence spanned a broader range of relaxation rates for more highly ordered bilayers like DMPC/Chol (Fig. 2B), and a narrower range for more fluid bilayers like DMPC (Fig. 2C), as seen in the experimental data (60,63).

**Figure 2.**
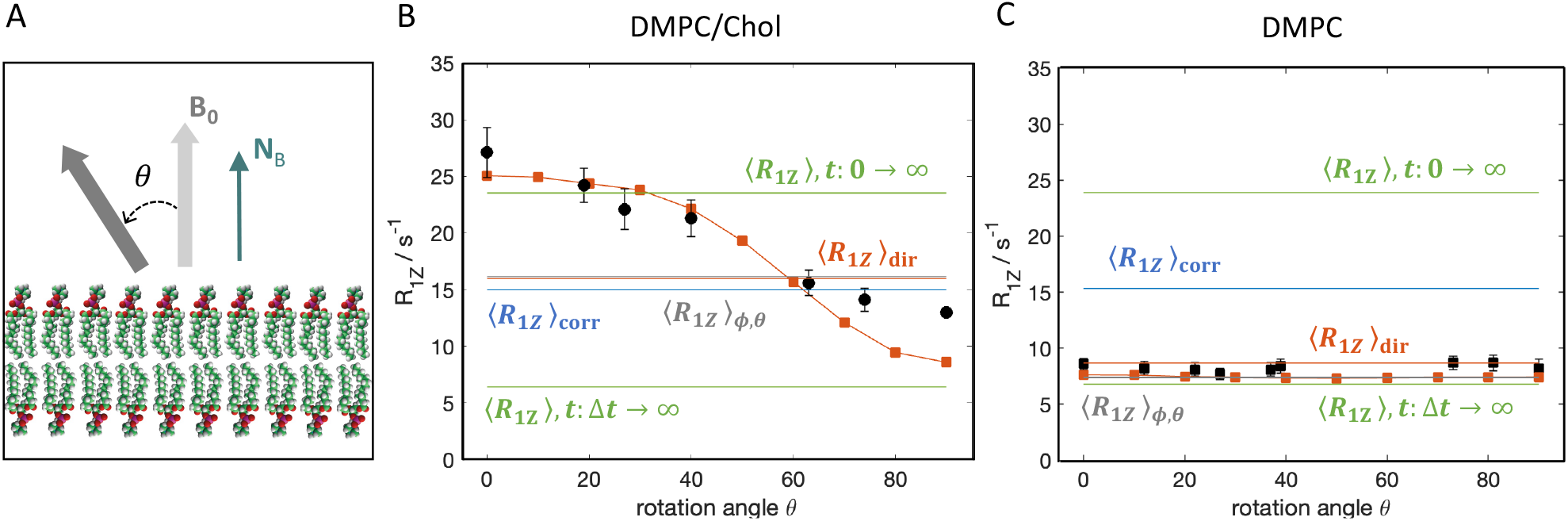
Angular dependence of CH bond dynamics in lipid simulations allows the calculation of orientationally averaged relaxation rates. (A) Schematic illustration of a bilayer in a simulation box. The magnetic field axis **B**_0_ is initially parallel to the bilayer director axis **N**_B_ (i.e., the *z*-dimension of the simulation box) so that the angle between them, *θ*, is 0 degrees. The **B**_0_ axis is then gradually rotated away from **N**_B_ by increasing the angle *θ*, and the CH bond relaxation rate is calculated with Eq. 6 in the laboratory frame, i.e., with respect to the **B**_0_ frame. (B) Calculated relaxation rate as a function of *θ* (red symbols) for carbon C9 on the *sn*-1 chain of DMPC in a bilayer with 50% cholesterol at 44°C. The orientationally averaged relaxation rate (grey) was calculated from the simulated angular dependence weighted by the uniform distribution of the directors on the surface of a sphere from Eq. 31. Shown for comparison are: experimental NMR data for carbons C7–C8 in oriented bilayers of DMPC/Chol 1:1 measured at 40°C from (60) (black symbols); the mean (green) and mean corrected (blue) orientation-independent relaxation rates; and the mean director-frame relaxation rate from Eq. 25 (red). (C) Calculated relaxation rate as a function of *θ* for carbon C12 on the *sn*-1 chain of DMPC in a single-component DMPC bilayer. Experimental NMR data are for carbon C13 in oriented DMPC bilayers measured at 40 °C from (63) and all colors and symbols are the same as in (B). **[2-column figure]**

Following the above protocol, the relaxation rates can thus be calculated relative to the fixed **B**_0_ laboratory frame for different angles *θ* without any averaging (Fig. 2). From the results, we estimated the orientationally averaged relaxation rate ⟨*R*_1Z_⟩_*θ,ϕ*_ by fitting the calculated angular dependence between 0 and 90 degrees, as well as its mirror image from 90 to 180 degrees to a smoothing spline function. Using the fit, we multiplied the results by *P*(*θ, ϕ*) from Eq. 31 and their sum yielded the mean value ⟨*R*_1Z_⟩_*θ,ϕ*_, which is plotted in Figs. 2B–C as a gray line. The ⟨*R*_1Z_⟩_*θ,ϕ*_ rate calculated for the DMPC/Chol bilayer (16.1 *s*^−1^) is clearly different from the average relaxation rate assuming constant probability at all *θ* (18.5 *s*^−1^), consistent with the dependence of *θ* on sin *θ* (Eq. 31). Together with the good agreement with experimental data, these observations thus further confirm the angular dependence of the CH bond fluctuations and demonstrate that simulations can provide a model-free estimate of the relaxation rate calculated directly in the lab frame.

### Angular anisotropy explains difference of orientation-independent and mean director-frame correlation functions

Notably, the orientationally averaged relaxation rates can also be approximated from numerical simulation data without explicitly considering the angular dependence of the relaxation. This is achieved by expressing the relaxation rate defined in the lab frame in Eq. 6, with the correlation functions and spectral densities in the director frame (Eq. 13), which are readily obtainable from MD simulations. We recently developed a framework that implements this approach by: (1) quantifying the CH bond orientations with respect to the bilayer normal (director) in Cartesian coordinates (Eqs. 1–3), (2) calculating their correlation functions (Eq. 22) and corresponding spectral densities in the director frame (Eq. 24), and (3) obtaining the *R*_1Z_ rates from the latter functions evaluated at the Larmor frequency *ω*_0_ and 2*ω*_0_ corresponding to the NMR measurement (Eq. 25). This approach assumes spherical averaging of the spectral densities in the lab frame 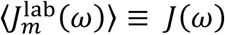 due to the uniform distribution of the bilayer directors (Eq. 12), but explicitly considers the three director-frame spectral densities 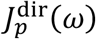 with respect to **N**_B_ that come from the ordering of the CH bonds relative to **N**_B_ (Eq. 8). We refer to the resulting MD-simulated values as the mean director-frame relaxation rates 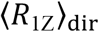, since they are quantified from the CH bond fluctuations relative to the director axis (bilayer normal). These calculated 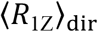 values, which are orientationally averaged by virtue of the averaged lab-frame spectral density *J*(*ω*) (Eq. 12), are plotted in Figs. 2B–C as horizontal red lines. For both bilayers with and without cholesterol, they are very similar to the corresponding orientationally averaged ⟨*R*_1Z_⟩_*θ,ϕ*_ relaxation rates.

For larger undulating membranes the calculation of the bilayer director **N**_B_ can be rather cumbersome, and an alternative approach circumvents the need for its explicit consideration. According to the well-established and widely used analysis of small molecules freely tumbling in solution, the model assumes that the measured relaxation rate does not have an angular dependence. This assumption simplifies the calculation significantly by allowing the application of the spherical harmonic addition theorem, which relates the orientationally averaged lab frame spectral density *J*(*ω*) from Eq. 12 to fluctuations of the angle 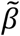 in Fig. 1C (Eq. 19). This angle quantifies the change in direction of the CH bond over time, rather than its orientation relative to a specific axis such as the **N**_B_ or **B**_0_ axes. The correlation function of the fluctuations, *C*(*t*), is expressed with the second-order Legendre polynomial of cos 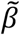 and is mathematically invariant to the rotation of the fixed axis as illustrated in Fig. 3 (64). We thus refer to the resulting relaxation rate ⟨*R*_1Z_⟩ as the orientation-independent relaxation rate (Eq. 20).

**Figure 3.**
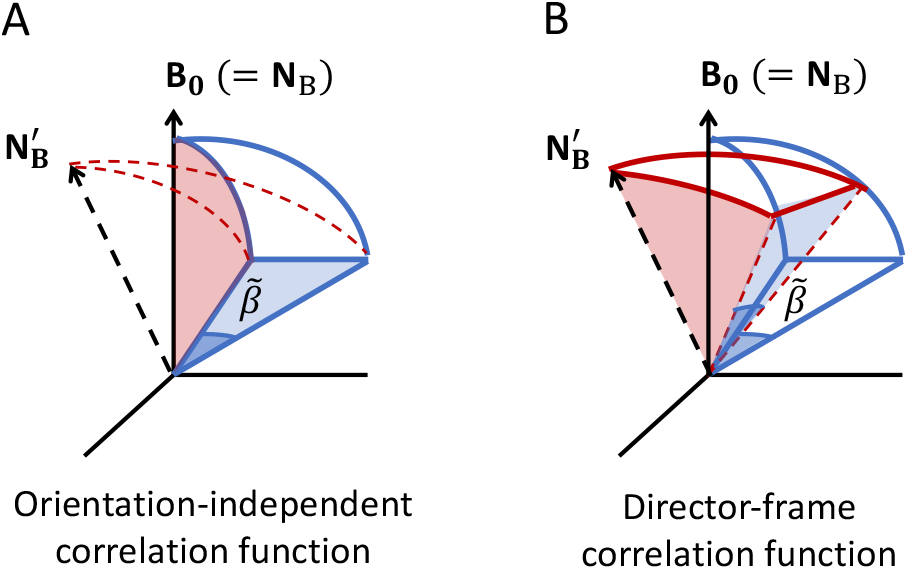
Illustration of why the spherical harmonic addition theorem is inapplicable to lipid membranes. The theorem considers the time-dependent angle 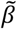 of the CH bond vector (solid blue lines) with itself and gives rise to the orientation-independent correlation function *C*(*t*). (A) Since there is no dependence on a specific fixed axis, 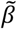 stays the same when the director **N**_B_ rotates, while the angles {solid blue, dashed red lines) it makes with the CH bond and the magnetic field axis **B**_0_ change. (B) In contrast, the director-frame correlation function *G*(*t*) is specific for the director axis and hence, rotation of **N**_B_ and the CH bonds {solid blue, dashed red lines) are *not independent*. Thus, as opposed to *C*(*t*), the *G*(*t*) function captures both the ordering potential characterized by **N**_B_ and the angular dependence of motion due to the director orientation **N**_B_ relative to the **B**_0_ frame. **[1-column figure]**

### Analysis of orientation-independent relaxation rates does not fully capture lipid dynamics

The spectral densities of CH bond fluctuations are two-sided Fourier transforms (FTs) whereas from simulations we can calculate FTs only as one-sided. The latter functions can be theoretically multiplied by a factor of two to yield the two-sided FTs (Eq. 26), from which the relaxation rate is then obtained (Eq. 27). Numerically however, this approach results in overcounting of the zeroth element of the correlation function which can artificially increase the result (see below). One approach to alleviate this problem is to completely disregard the zeroth (*k* = 0) element, i.e., to perform a Fourier transform of the correlation function *C*(*t*) for *t* ≥ *Δt*, which will introduce a frequency-dependent phase shift of the spectral density. An alternative approach is to include the zeroth element, but to ensure it is not overcounted (Eq. 28) to yield what we will call the mean *corrected* ⟨*R*_1Z_⟩_corr_ orientation-independent relaxation rate.

The orientation-independent relaxation rates ⟨*R*_1Z_⟩ calculated for *t* ≥ 0 and for *t* ≥ *Δt* are plotted in Figs. 2B–C as horizontal green lines. They are quite different from each other, and even though one of them overlaps almost perfectly with ⟨*R*_1Z_⟩_*θ,ϕ*_ in the DMPC bilayer (Fig. 2C), neither falls inside the range defined by the angular dependence of *R*_1Z_ for both the DMPC/Chol and DMPC bilayers (Figs. 2B–C). This is not surprising considering the previously observed sensitivity of the results to the incorporation of the zeroth (*k* = 0) element of the *C*(*t*) orientation-independent correlation function (23). In comparison, the corrected mean orientation-independent relaxation rate ⟨*R*_1Z_⟩_corr_ (horizontal blue lines) has an intermediate value, and is closer to the orientationally averaged ⟨*R*_1Z_⟩_*θ,ϕ*_ rate and the mean director-frame relaxation rate ⟨*R*_1Z_⟩_dir_ in DMPC/Chol (Fig. 2B) but still outside of the angular range for the DMPC bilayer (Fig. 2C). Here the main difference between the orientation-independent relaxation rates ⟨*R*_1Z_⟩ and ⟨*R*_1Z_⟩_corr_ and the orientationally averaged ⟨*R*_1Z_⟩_*θ,ϕ*_ relaxation rate is the assumption of the spherical harmonic addition theorem with no angular dependence. Thus, the inability of ⟨*R*_1Z_⟩ or ⟨*R*_1Z_⟩_corr_ to accurately quantify the dynamics of the CH bonds with and without cholesterol can be explained by the different mobility and order of the bilayers. When the lipid chains are more disordered as in DMPC, the CH bonds sample a wider range of angles with respect to the bilayer director in contrast to DMPC/Chol, where they are more ordered and exhibit restrained motions within a confined orientation. These motional regimes affect the diffusion of the lipids in the bilayer and the angular dependence of the relaxation, which is either more enhanced or suppressed. Consequently, the approximation of the CH bond dynamics by monitoring only changes in 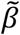 fails to capture this full spectrum of molecular dynamics.

### Output frequency of atomic coordinates affects accuracy of calculated relaxation rates

In simulations the relaxation rates are obtained from the spectral densities of the CH bond fluctuations (Eqs. 6, 13, 20) near the resonance (Larmor) frequency *ω*_0_ and twice this value. Regardless of how the fluctuations are quantified, e.g., in Cartesian coordinates with Wigner *D*-functions as in Eqs. 1–3 or with the angle 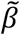 and spherical harmonics as in Eq. 19, the spectral density is calculated from the Fourier transform of the respective autocorrelation function. In theory, that should be a two-sided continuous Fourier transform (or CFT) [see Eq. 11 in (23)]. However, simulations are run with a discrete time step *Δt*_sim_ and atomic coordinates are output at some fixed sampling time interval *Δt* ≥ *Δt*_sim_, making the simulation data points spaced *Δt* time units apart. Consequently, the calculated spectral density is inherently a one-sided and discrete Fourier transform (or DFT). A related aspect is that for a correlation function with a sampling time interval of *Δt* there is an upper limit to the frequency that can be detected. Beyond this limit, positive difference frequencies cannot be distinguished from negative difference frequencies. Thus, in the Fourier transform all frequencies greater than 1/2*Δt* are *backfolded* onto the spectral density, i.e., frequencies of 1/2*Δt* + υ cannot be distinguished from frequencies of 1/2*Δt* − υ, contributing to the offset (Eq. 24). The highest frequency in the DFT that can be sampled is given by 1/2*Δt* according to the Nyquist-Shannon sampling theorem.

A one-sided DFT can then be used to approximate the two-sided CFT by multiplying the former by a factor of two. Yet, special care must be taken not to overcount the element at the lag time *k* = 0, which is the largest element of the autocorrelation function. That is why the director-frame spectral densities in Eq. 24 and *J*_c_(*ω*) in Eq. 28 are written as the sum of the Fourier transform of the correlation function at *k* = 0 and twice the Fourier transform of the correlation function for *k* ≥ 1. For a generalized spectral density *J*_gen_(*ω*) and correlation function *G*_gen_(*k*), this summation reads follows:

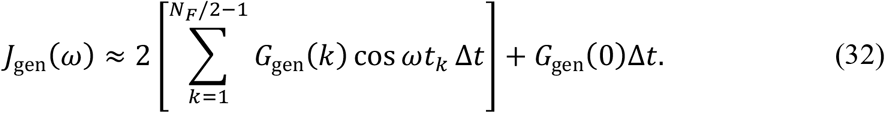

Since *G*_gen_(0) is equal to the variance of the fluctuations for *G*(*t*) (Eq. 23), or 1 for *C*(*t*), the second term in Eq. 32 is independent of the frequency *ω* and represents a constant term proportional to the sampling time interval *Δt*. This constant is added to the spectral density at every *ω* giving a frequency-independent offset. In theory, when *Δt* → 0 this term becomes infinitesimally small. Yet, in simulations the choice of *Δt* is bounded below by the simulation time step *Δt*_sim_ (usually 2 fs for fully atomistic simulations) but is often orders of magnitude larger, *Δt* ≫ *Δt*_sim_, e.g., pico- or nanoseconds, and is somewhat arbitrary. This can introduce an artificial shift in the spectral density across all frequencies and directly affect the accuracy of the calculated relaxation rates.

Figure 4 shows this problem for the mean director-frame relaxation rate ⟨*R*_1Z_⟩_dir_ (left), as well as for the corrected orientation-independent relaxation rate ⟨*R*_1Z_⟩_corr_ (right). In both cases there is an apparent increase in the calculated relaxation rate as *Δt* increases. However, while for ⟨*R*_1Z_⟩_corr_ the change is a constant offset as expected from Eq. 28, the differences in the ⟨*R*_1Z_⟩_dir_ values vary along the chain and are consistent with the respective dynamics of the chain segments. For instance, the CH bonds closer to the bilayer midplane which are more disordered and have shorter correlation times (see below) show greater sensitivity as *Δt* is increased from 4 ps to 40 ps and 80 ps, while the CH bonds closer to the headgroup whose correlation times are on the order of 30 ps (see below), begin to increase only when *Δt* goes up to 80 ps.

**Figure 4.**
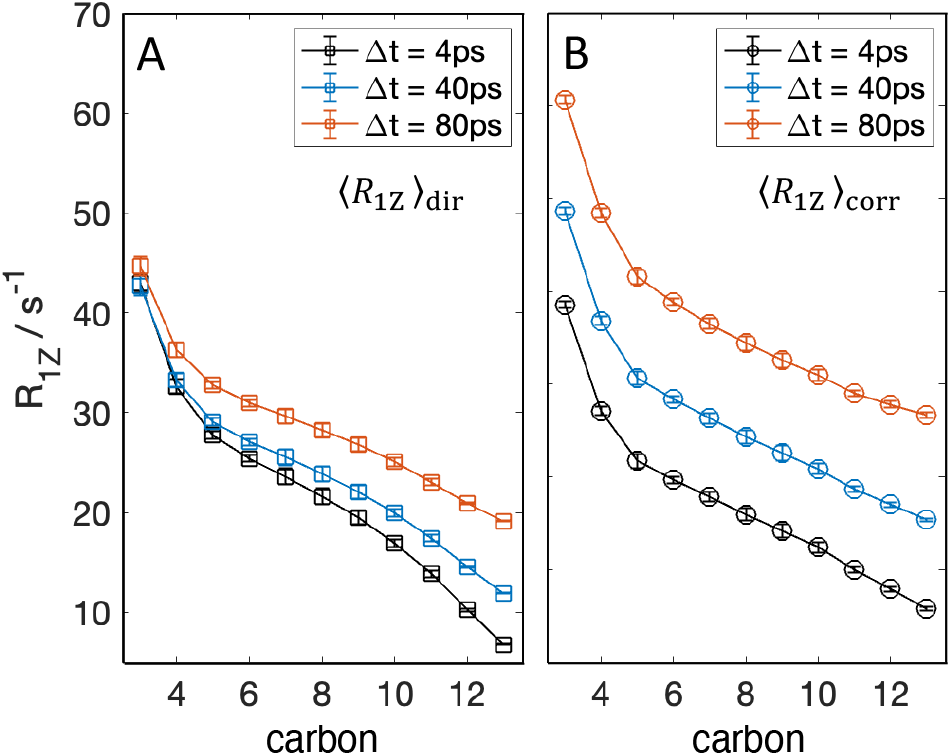
Calculated relaxation rates depend on the output frequency of atomic coordinates. (A) Mean director-frame relaxation rates and (B) corrected orientation-independent relaxation rates calculated from the same 800-ns trajectory of a DMPC bilayer with 50 mol% cholesterol but with different output frequencies of atomic coordinates. The simulation was run with a 2-fs timestep and data for analysis were taken every *Δt* ps. An apparent increase in the relaxation is observed as the temporal resolution is lowered due to the discrete calculation of the Fourier transform of the fluctuations’ autocorrelation function. All simulations were performed at 44°C. **[1-column figure]**

### Carbon–hydrogen bonds exhibit both slow and fast dynamics in lipid bilayers

According to the Nyquist-Shannon sampling theorem, the discrete Fourier transform in Eq. 32 can accurately represent the underlying continuous Fourier transform if *Δt* is smaller than half the period of the fastest CH bond motion in the bilayer. To determine the range of CH bond motions in a bilayer, we first analyzed the autocorrelation function of the fluctuations in 2-*μ*s-long simulation trajectories of DMPC without and with 33 mol% cholesterol from Ref. (23). Following the approach of Lipari and Szabo (49), we calculated the effective correlation time, τ_eff_, of the CH bond reorientations at all carbons on the *sn*-2 chain of DMPC using Padé approximants (see Eq. 30 in Methods). Figure 5A shows the results for the bilayer without cholesterol, where τ_eff_ was between 0.5 and 1 ns, while in the bilayer with cholesterol τ_eff_ reached 4 ns, consistent with the slower dynamics and higher ordering of the CH bonds due to the sterol. For these simulations, output coordinates were saved every *Δt* = 40 ps, which according to Fig. 5A is smaller than one-half of τ_eff_, that is *Δt* < (1/2)τ_eff_, across all carbons.

**Figure 5.**
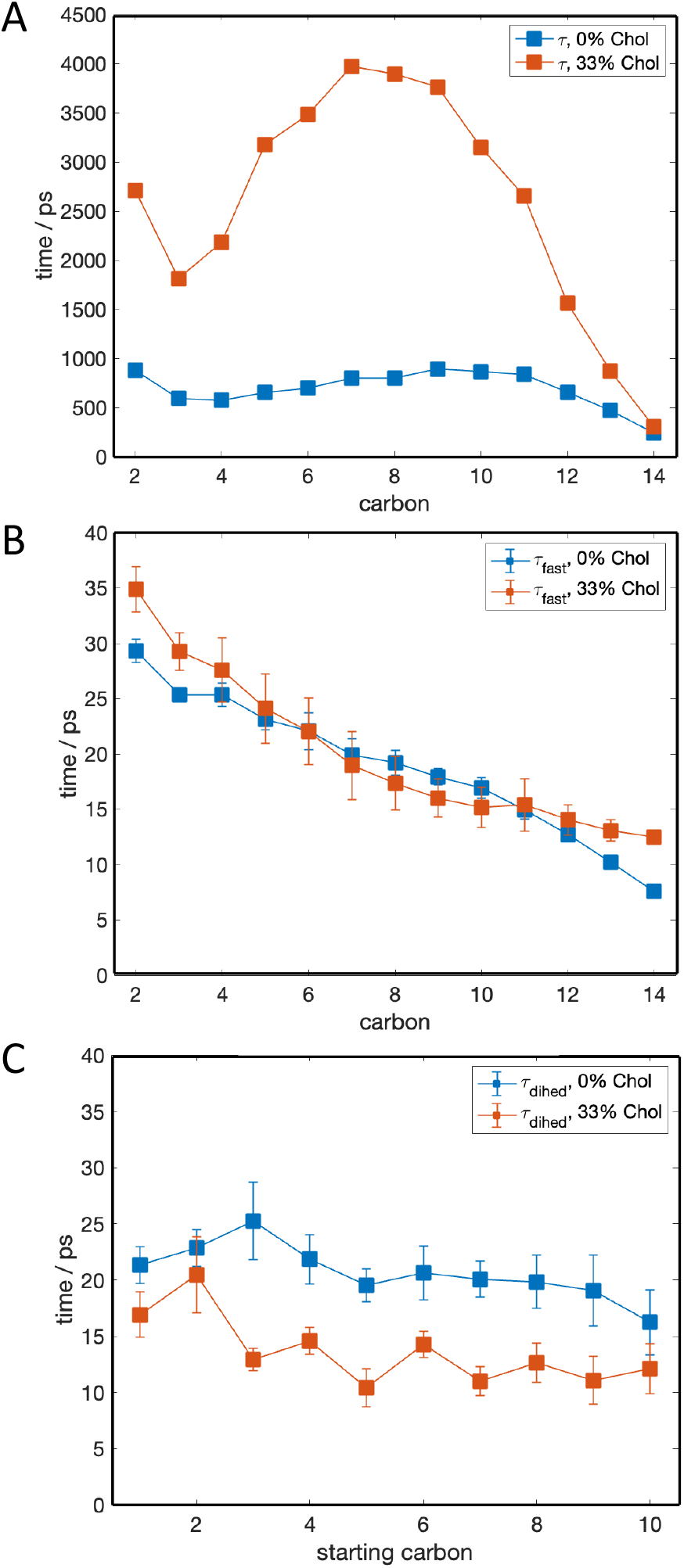
Carbon–hydrogen (CH) bond dynamics of lipid membranes have both slow and fast relaxations. (A) Effective correlation time of CH bond fluctuations calculated from 2-*μ*s-long trajectories of fluid DMPC bilayers with 0 and 33 mol% cholesterol. Atomic coordinates were output every 40 ps. (B) Effective correlation time of CH bond fluctuations calculated from short 400-ps-resampling simulations with output time interval of 10 fs. (C) Isomerization times of carbon segments along the *sn*-2 chain of DMPC calculated from the short resampling simulations from (B). Shown are the correlation times of *trans*-to-*gauche* and *gauche*-to-*trans* isomerizations of four consecutive carbon segments quantified from changes in their dihedral angles. All simulations were performed at 44°C. **[1-column figure]**

To examine the potential contributions of faster dynamics of the CH bonds, we resampled the long simulations by running multiple short simulations. Here we chose a value of *Δt*_rs_ = 10 fs instead, starting at different time points of the original trajectories as described in Methods. Calculation of τ_eff_ from these resampling simulations at higher temporal resolution showed a much faster reorientation of the CH bonds, τ_fast_, occurring on a sub-40-ps timescale (Fig. 5B). These fast correlation times ranged between 5 and 35 ps and were independent of cholesterol concentration. This observation is consistent with results from ^13^C relaxation time experiments which are sensitive to fast motions (high frequencies) and reveal no significant effect of cholesterol on the local fast lipid dynamics (46). Importantly, their existence makes the above analysis inconsistent with the Nyquist-Shannon sampling theorem with standard choices of *Δt* ≫ *Δt*_rs_, including the 40-ps output interval employed in the original long trajectories from Ref. (23).

Next, we further investigated the origins of the fast CH bond correlation times by analyzing the isomerization rates of the carbons along the acyl chains (Fig. 5C). Every four consecutive carbons along a lipid chain can exist in either *trans* or *gauche* isomers as defined by their dihedral angle. The rate with which each segment changes its isomerization has been shown to be very fast in alkanes, having correlation times of ∼20 ps (65,66). While these calculations have caveats, e.g., they are model-based and do not consider the concerted transitions between conformers (67), they provide an estimate of the timescale of the dynamics. We therefore followed the approach introduced by Richard Pastor et al. for analysis of isomerization rates of peptide chains (Eqs. 5–7 in (54)) to estimate the isomerization rates of the lipid chain segments in our simulated bilayers. We classified the instantaneous conformation of a segment as *trans* or *gauche* based on its dihedral angle and used the resulting time series to calculate a number correlation function *C*_N_(*t*), as described in Methods. The corresponding effective correlation times τ_dihed_, calculated from *C*_N_(*t*) with Eq. 30, are plotted in Fig. 5C. For both bilayers, with and without cholesterol, the isomerization rates were very similar to the respective τ_fast_ values in Fig. 5B, indicating that the cholesterol-independent fast correlation times of the CH bond motions likely arise from the rapid isomerizations of the chain segments. Taken as a whole, our results confirm that the CH bond dynamics span multiple timescales that are orders of magnitude apart. This characteristic feature of the fluctuations has consequences for any type of multiscale analysis that relies on proper sampling of the fastest motions.

### Multiscale simulations can overcome the Nyquist limit by resampling carbon–hydrogen bond fluctuations

The existence of fast CH bond dynamics presents both a challenge and an opportunity for the accurate calculation of a continuous Fourier transform from multiscale discrete simulation data (Eq. 32). This is due to the relatively large time interval employed for practical outputting of the atomic coordinates, especially in long trajectories, which prevents adequate sampling of the dynamics of the system according to the Nyquist-Shannon theorem. In other words, the data points of the autocorrelation function of the fluctuations from Eq. 22 are spaced too far apart, making the last constant term on the right in Eq. 32 (also Eqs. 24 and 28) too large and introducing an artificial shift in the spectral density (Fig. 4). One way to alleviate this problem is to reduce *Δt* as we did in the 400-ps resampling simulations utilizing *Δt*_rs_ of 10 fs. However, the poor sampling in each individual trajectory limits the accessible frequencies and makes the accurate calculation of relaxation rates challenging.

Alternatively, we have shown that the autocorrelation function 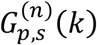 of the lipid CH bond fluctuations, quantified by the Euler angles *β* and *γ* and the Wigner rotation matrix elements from Eqs. 1–3, follows a simple mathematical expression (23). In particular, it can be fit to a power-law of the form *ax*^c^ + *c* as shown in Fig. 2A in (23). This is important because it allows us to use the best fit 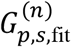 to resample the correlation function at a much smaller *Δt*_dDe_ ≪ *Δt*. Since 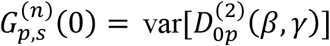 is the largest element of the autocorrelation function (ACF), we can perform the fit on 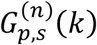 for *k* ≥ 1 and find the smallest *Δt*_fit_ that allows the ACF to smoothly approach the variance of the data as *k* → 0 as shown in Fig. 6A. For that, we start from *Δt*_fit_ of 100 fs and increment it by 100 fs until at lag *k* = 1 (or equivalently, *t* = *Δt*_fit_), the value of 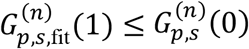. This approach yields *Δt*_fit_ values between 0 and 30 ps, which are one-half or less than one-half of the corresponding fast correlation times τ_fast_ for most carbons (Fig. 6B). The recovered spectral density then reads:

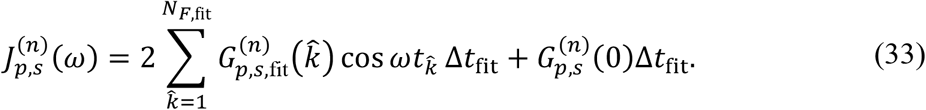

According to the Nyquist-Shannon theorem, it accurately describes the two-sided continuous Fourier transform (CFT) of the correlation function, thus alleviating the sampling problem introduced by the large output interval of atomic coordinates. Calculating the spectral density with Eq. 33 produced relaxation rates in excellent agreement with experiment using 2-*μ*s simulations with output frequency *Δt* of 40 ps for DMPC bilayers with increasing amounts of cholesterol (23).

**Figure 6.**
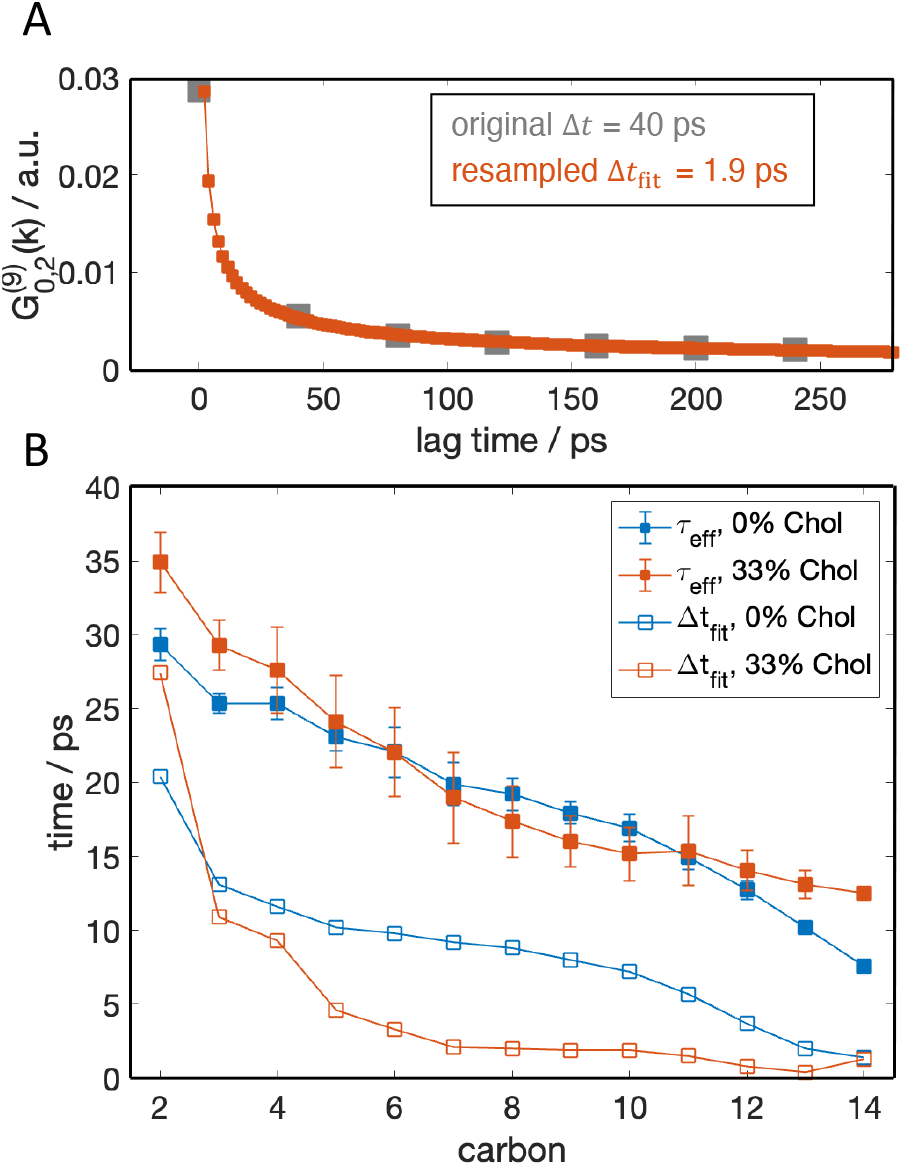
Mathematical resampling of CH bond autocorrelation function bypasses sampling limit in discrete Fourier transformation for lipid membranes. (A) The autocorrelation function of the CH bonds at carbon C9 on the *sn*-2 chain of DMPC in the bilayer with 33 mol% cholesterol, 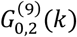, calculated every 40 ps from the original long trajectories (gray) and resampled every 1.9 ps from the best power-law fit to 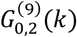 (red). (B) The smallest resampling time interval *Δt*_fit_ obtained from best fits to autocorrelation functions 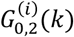 for carbons 2 ≤ *i* ≤ 14 in the two simulations (open symbols). Shown for comparison are fast correlation times τ_fast_ for the corresponding CH bonds calculated from short resampling simulations with output time interval of 10 fs and replotted from Fig. 2B. All simulations were performed at 44°C. **[1-column figure]**

To examine in more detail the accuracy of our estimate obtained using Eq. 33, we extended the original 2-*μ*s trajectories of DMPC and DMPC/Chol by an additional 800 ns, during which the atomic coordinates were output every 4 ps. According to the results in Fig. 5B, a *Δt* of 4 ps should meet the criteria of the Nyquist-Shannon theorem, and allow for direct calculation of the relaxation rates without having to first fit the correlation function, i.e., the spectral density can be obtained from Eq. 24 instead of Eq. 33. Figure 7 shows a comparison between the mean director-frame relaxation rates calculated with Eq. 25 by using either *Δt* of 4 or 40 ps without fitting the correlation function, or *Δt* = *Δt*_fit_ from Fig. 6B obtained by fitting the correlation function from the raw data output every 40 ps. As seen in both bilayers, fitting the correlation function successfully removes the artificial increase in the relaxation rates introduced by the larger output frequency of 40 ps, and recovers almost perfectly the results obtained when *Δt* is 4 ps. Thus, this approach represents an effective solution that can be used to calculate reliable estimates of the mean director-frame relaxation rates of CH bonds from data with lower temporal resolution. Since the orientation-independent *C*(*t*) function does not follow a power-law, such mathematical resampling cannot be applied to alleviate the *Δt*-dependence of the mean ⟨*R*_1Z_⟩ and mean corrected ⟨*R*_1Z_⟩_corr_ orientation-independent relaxation rates (Fig. 4B).

**Figure 7.**
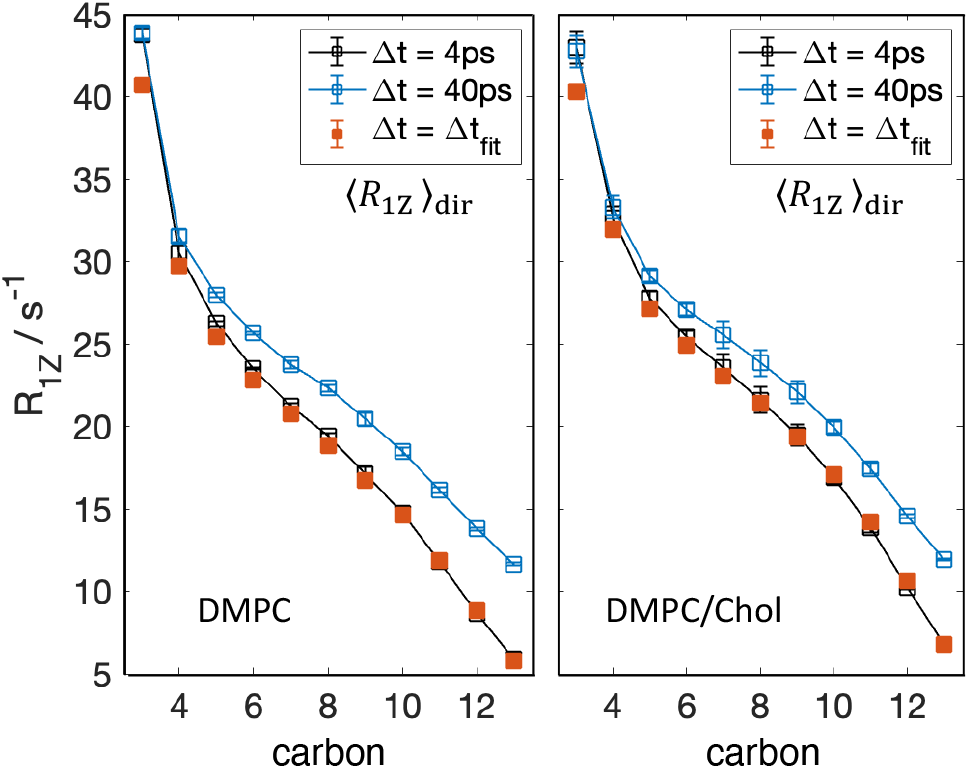
Fitting the correlation function of bond fluctuations produces reliable estimates of their relaxation rates. (A) The CH bond relaxation rates calculated with Eq. 25 from simulation data with 40-ps (blue) or 4-ps (black) resolution without fitting the correlation function show differences due to the dependence on *Δt* (cf. Fig. 4A). Fitting the correlation function of the bond fluctuations from the low-resolution data and resampling it at a much smaller *Δt*_fit_ (red) alleviates this problem and recovers the results from the higher resolution data. (B) The plotted mean director-frame relaxation rates are for the carbons on the *sn*-2 chain of DMPC in the bilayer without cholesterol (left) or with 33 mol% cholesterol (right). All simulations were performed at 44°C. **[1-column figure]**

## DISCUSSION

Biomolecular simulations are uniquely informative in detecting the functional motions of lipids in membranes together with complementary theoretical and experimental biophysical methods (1,22,29,68-78). Theoretical simulations in particular have attracted recent notice with the advent of artificial intelligence (AI) approaches in addition to physics-based force fields (79). Such techniques are expected to contribute strongly to future data-driven methods in structural and cellular biology. As an ideal benchmarking technique, solid-state NMR delivers experimental order parameters for the carbon–hydrogen (CH) bond fluctuations along the lipid chains, while the relaxation rates access the corresponding dynamics. However, while actual CH bond reorientations occur on pico- and nanosecond timescales, their NMR relaxation rates at the Larmor frequencies of conventional instruments are in the millisecond regime (44,45,63,80,81). Relatively short multiscale simulations allow the study of femto-, pico-, and nanosecond dynamics, which can be used to quantify longer millisecond-time relaxation rates by Fourier transformation of the autocorrelation function of the CH bond fluctuations. To accomplish this goal, the liposome geometry in an actual NMR sample (62) requires evaluation of the orientational averaging of the simulated dynamics obtained from flat bilayer patches over the longer experimental time scale. Here we show that such averaging can efficiently be performed while accounting for the angular dependence of the relaxation, which is critical for validating results for membranes with different mobility. Present computational approaches yield an efficient framework for numerical calculations but are unable to capture these crucial aspects of the orientational dependence of the bond dynamics in lipid membranes.

We previously developed a robust framework that enables this calculation to be efficiently carried out and allows for the simulation results to be directly compared to experimental NMR values (23). In the process, we discovered a strong dependency of the calculated relaxation rates on the time resolution of the simulation data points, i.e., the output interval of the trajectory frames used for the analysis. We hypothesized that the issue was due to the presence of very fast dynamics of the CH bonds, which prevented the accurate calculation of a continuous Fourier transform with the employed output interval of 40 ps. Accordingly, we directly tested this hypothesis by resampling the simulation trajectories at different time points with short 400-ps-long simulations and an output frequency of 10 fs. The new high-resolution data confirmed the presence of CH bond fluctuations with correlation times between 5 and 35 ps, as observed previously by Pastor and coworkers for a DPPC bilayer (66). Further analysis indicated that the hierarchical dynamics are mediated by the changes in isomerization rates of the carbons along the lipid chains. A resampling approach presents a solution to the data problem by using a fit to the correlation function to satisfy the conditions of the Nyquist-Shannon sampling theorem (23). In the present study we directly evaluated the accuracy of this mathematical resampling, showing that it successfully recovers the relaxation rates obtained from high-resolution data (Fig. 7). Our results reveal the presence and source of the very fast CH bond dynamics and further establish the validity of resampling their correlation functions, thus increasing the effective time resolution.

### Explicit consideration of bilayer director is essential for calculating relaxation rates

Differences in the CH bond relaxation rates obtained with the classical status quo approach and with our recently developed computational framework naturally raise questions about the validity of the underlying assumptions. In particular, the theory behind the commonly applied methodology comes from solution NMR (50,82) and can be traced back to application of the spherical harmonic addition theorem (Eq. 18) as in the rotational relaxation of isotropic liquids. This theorem assumes that there is no fixed alignment axis relative to which the CH bond fluctuations are calculated (Fig. 3) and thus produces orientation-independent relaxation rates as in textbooks. As such, the results are unable to account for the effects of the anisotropic relaxation observed both experimentally and computationally, or to accurately describe lipid motions in both more fluid and more ordered environments (Fig. 2). It is unavoidable that relating simulations and experiments requires averaging of the simulation results to achieve correspondence with the rates measured from liposome samples with solid-state NMR spectroscopy. Such averaging can be carried out either at the level of the correlation function (Eq. 14) or spectral density (Eq. 12) and produce equivalent results. One possible misconception is that because the orientation-independent correlation function *C*(*t*) and the mean director-frame correlation function *G*(*t*) are both spherically averaged they are identical apart from a multiplicative factor (Eq. 18). Formally the mean director-frame correlation function *G*(*t*) is related to the orientation-independent *C*(*t*) correlation function by *G*(*t*) = (1/5)*C*(*t*), but that equivalence holds *only* under the assumption of the spherical harmonic addition theorem that there is no fixed director axis. Still, the existence of the CH bond order parameters clearly illustrates that there is a potential of mean force (PMF) or ordering induced on the lipids due to an alignment frame characterized by the director (Fig. 3) (56). By contrast, the use of *C*(*t*) bypasses the dependence on a director and instead considers the self-correlation of the CH bond fluctuations. This treatment means that isotropic motion is implicitly assumed as in the case of the Debye model for rotational relaxation in liquids (83). Comparison of the results with solid-state NMR data (Fig. 2) shows that explicit consideration of a fixed director axis, which is the main difference between the two approaches, is essential for the accurate estimation of the experimentally measured relaxation rates.

### Multiscale simulations of biomembranes are limited by output frequency of atomic coordinates

In recent years computational resources have become more easily accessible by researchers and educators, as in the case of graphics processing units (GPUs) used in computer game applications. This has been facilitated by the establishment of large supercomputing infrastructures made available to the public via open proposal calls and freely available training programs on their use (84). As a result, microsecond-long simulations have become standard even for relatively large fully atomistic systems (85,86). The decision of how often to output atomic coordinates during a simulation determines both the amount of space needed to store the trajectory files and the maximum time resolution of any subsequent analyses. For instance, an all-atom bilayer with 200 lipids and 45 water molecules per lipid amounts to about 50,000 atoms. Simulating this system for 10 *μ*s would require roughly 200 GB of storage if coordinates are output every 40 ps and 200 TB of disk space if the data output *Δt* is 40 fs. Therefore, the time resolution needs to be balanced with practical considerations, where simultaneous access to sub-picosecond and microsecond dynamics is often challenging for multiscale membrane dynamics (22,76,85,87-89). Even so, relatively small windows at faster dynamics can be created by resampling long trajectories via short simulations with smaller sampling intervals *Δt*_rs_ < *Δt* (90). While this approach provides access to faster molecular motions, the statistics are limited, and the results are strongly dependent on the state of the bilayer at the chosen starting points for resampling. Lipids can take ∼100 ns to exchange places with their nearest neighbor and tens of hundreds of nanoseconds to equilibrate their lateral distribution, even in a simple binary mixture (91). Thus, on the timescale of 400 ps for example, corresponding to the short resampling simulations we performed here, the overall lipid organization in the bilayer remains the same. Reducing *Δt* by an order of magnitude as a way to increase the resolution of the CH bond autocorrelation function and facilitate the accurate calculation of its Fourier transform is often not feasible. Here, we show that relatively long simulations (of 800 ns) with an output interval of 4 ps can provide a reasonable solution.

### Mathematical resampling based on power-law decay gives solution to the data problem

To increase the effective sampling of the simulation data, we take advantage of the shape of the CH bond autocorrelation function calculated in the director frame with Eq. 21 (Fig. 1). When the orientation of the CH bonds is defined with the Wigner rotation matrix elements from Eqs. 1–3, the autocorrelation function (ACF) of their fluctuations (Eq. 7) follows a power-law function (23,45,51). Fitting the ACF to a power-law then allows us to sample it at a much smaller *Δt*_fit_ ≪ *Δt*, bypassing the dependence of its Fourier transform on the arbitrary choice of *Δt* to avoid aliasing. While the identified *Δt*_fit_ values are bounded below by the smooth transition of the ACF fit to the element at lag time *t* = 0, i.e., the variance of the fluctuations (Eq. 23), they are still small enough to satisfy the Nyquist-Shannon sampling theorem. This eliminates the artificial *Δt*-dependent shift in the spectral density (Eq. 32), producing relaxation rates in excellent agreement with those calculated from simulation data with higher resolution (Fig. 6), as well as those directly obtained from NMR experiments (Fig. 6 in (23)). Searching for the smallest *Δt*_fit_ to accurately calculate relaxation rates from discrete simulation data appears to be the counterpart of extrapolating experimental spin-lattice *R*_1Z_ relaxation rates to infinite frequency to evaluate the local CH bond fluctuations (Fig. 6, see below) (52,66,92). In the present approach, we quantified the CH bond orientation with respect to the bilayer normal (director axis) with the Euler angles *β* and *γ* in Cartesian space, but this can also be done with the spherical angles *θ* and *ϕ* in a similar way (Fig. 1C). Either representation necessitates orientational averaging to facilitate comparison with NMR data from liposome dispersions of lipids. In the status quo approach this averaging is done on the correlation function, as discussed above. However, the resulting averaged correlation function *C*(*t*) is equal to 1 at lag time 0 and does not follow a simple functional form. Notably, this prevents application of the mathematical resampling approach, which relies on a fit to the correlation function. Consequently, the resulting orientation-independent relaxation rates have a strong dependency on the output interval in the trajectories as shown in Fig. 4, and fail to capture the full spectrum of the CH bond dynamics regardless of the precise implementation of the discrete Fourier transformation of the *C*(*t*) correlation function (Fig. 2).

### Data-driven simulations reveal collective lipid motions due to acyl chain isomerizations

Further consideration indicates that the source of the sampling problem in the Fourier transformation of the CH bond correlation function is the presence of very fast reorientations of the bonds relative to the time resolution of the simulation data points. Evidently they have correlation times of less than 40 ps (Fig. 5B) and become faster towards the ends of the chains at the bilayer midplane, in agreement with observations made 30 years ago by Richard Pastor et al. from simulations shorter than our resampling trajectories (66), consistent with NMR findings (92). Additionally, we find these fast motions are independent of cholesterol concentration in the membrane, implying they are not influenced by bulk or mesoscale bilayer properties. In contrast, the CH bonds also exhibit slower dynamics with correlation times that are orders of magnitude larger than 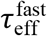 and thus are more easily discoverable with current standard choices for the time step of outputting atomic coordinates (51). These slower dynamics have a more complicated dependence on chain carbon depth within the bilayer and are strongly influenced by the ordering effects of cholesterol (15).

Building on earlier research (66), the ability to access logarithmic time-scales in our simulations via the resampling approach allows us to analyze the origins of the fast CH bond dynamics. Our analysis indicates that these fast dynamics are likely related to the rapid changes in isomerization of the carbon segments along the lipid chains (Fig. 5C). This proposal can explain why the resulting correlation times are the same within error for bilayers with and without cholesterol, i.e., they are not affected by the structural and dynamical effects induced on the lipids by the sterol. To further confirm the universality of the fast CH bond dynamics, we can analyze the respective changes in relaxation rates as a function of inverse frequency (Fig. 8). Being a characteristic signature of the spectral density, the relaxation rate at a given frequency (*ω*) describes the extent of correlation of the bond fluctuations on that timescale (*t* ≈ 1/*ω*). At all carbons on the lipid chains, both in the presence and absence of cholesterol, the CH bond relaxation rates differ at relatively small frequencies (or larger *ω*^−1/2)^ power law as *t* → ∞). Yet they converge at infinite frequency as *t* → 0 indicating that the underlying fast motions of the bonds are similarly correlated across all datasets at very short timescales (Fig. 8). In fact, NMR analysis (44,92-94) shows that this common relaxation rate at infinite frequency is the same as that of simple liquid hydrocarbons in solution (not in a bilayer), suggesting that the fast dynamics are a manifestation of the local bilayer microviscosity, coming from the hydrocarbon environment (44,66). The same conclusion was made from direct comparison of the fast CH bond dynamics in a 1,2-dipalmitoyl-*sn*-glycero3-phosphocholine (DPPC) bilayer to those of neat hexadecanes, i.e., hydrocarbons of the same length as the lipid chains but in solution instead of a bilayer (93). Towards the ends of the lipid chains, the CH bonds in the DPPC bilayer have the same correlation times as those of the equivalent carbon positions in hexadecane. However, the CH bonds at the top of the DPPC chains move much slower, suggesting that the dynamics in that region are strongly influenced by the interfacial tension at the bilayer surface. These observations indicate that the CH bonds of lipids in a bilayer do indeed experience the same microviscosity as liquid hydrocarbons, but only when sufficiently far from the bilayer/water interface.

**Figure 8.**
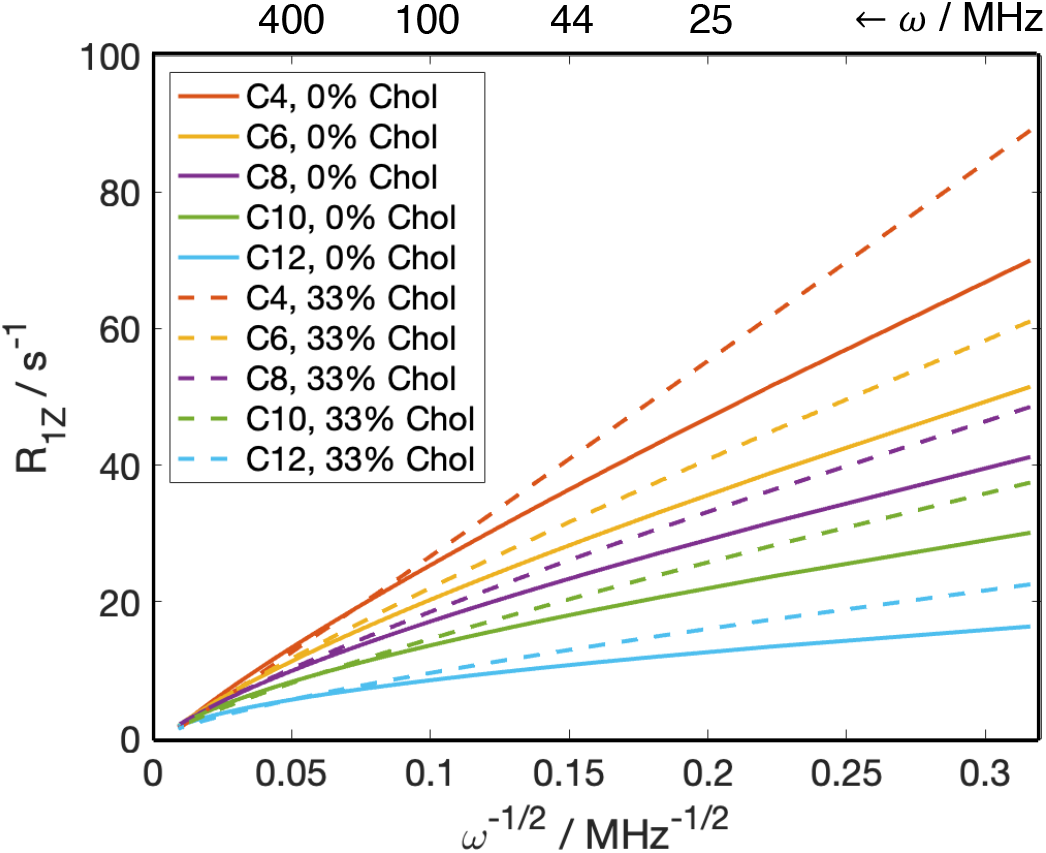
Fast carbon–hydrogen fluctuations are universal descriptors of bilayer microviscosity. Calculated mean director-frame relaxation rates of CH bonds at carbons 4, 6, 8, 10, and 12 on the *sn*-2 chain of DMPC in bilayers with 0 and 33 mol% cholesterol are plotted against inverse square root of the Larmor frequency *ω*. Irrespective of lipid composition, the relaxation rates at all carbons converge at infinite frequency (or infinitesimally small lag time). Hence the fast correlation times of the CH bonds are universal descriptors of the bilayer hydrocarbon environment corresponding to the local microviscosity. **[1-column figure]**

### Simulated lipid dynamics are validated with solid-state NMR spectroscopy

As mentioned above, our new and improved framework for the calculation of NMR relaxation rates from simulations addresses various issues that have come up in physics-based MD simulations of lipid membranes. While the absolute relaxation rates of carbons in the simulations and experiments are directly comparable, further validation can be achieved against the relationship between CH bond relaxation rates and their respective squared order parameters (44,92,93). We found that the two quantities exhibit the so-called square-law, which is clearly discernible both in the experimental (95,96) and simulation data (23). This dependence describes the nature of lipid segmental dynamics resembling those of nematic liquid crystals. We also found that it holds specifically for the CH bonds situated far from the lipid/water interface where the two leaflets intercalate [see Fig. 5 in (23)]. The unique resemblance of the microviscosity of this bilayer region to that of neat hydrocarbons whose relaxation rates are independent of frequency (Fig. 7) implies that the square-law relationship is an intrinsic property of the bilayer core. Furthermore, the slope of the square-law uniquely corresponds to the local bilayer bending rigidity, thus revealing packing trends in the elastic behavior of lipid bilayers that are sometimes concealed when measured at long length- and time-scales (15,97,98). Therefore, validating lipid dynamics with NMR spectroscopy through the square-law dependence provides an unprecedented opportunity to corroborate multiple aspects of membrane biophysical properties captured by the underlying lipid force fields.

## CONCLUSION

Molecular dynamics simulations offer a fascinating and detailed look into the structural and dynamical properties of biological systems. However, the generality and reliability of the results depends on how well the simulations capture the essential aspects of the processes they model. Being sensitive to the fluctuations of individual carbon–hydrogen bonds in a bilayer, NMR relaxation provides different points for comparison and validation of the membrane dynamics. Extracting comparable relaxation rates from the simulation trajectories is not trivial, as it relies on underlying theoretical concepts and continuous Fourier transformation of discrete simulation data. Our new framework successfully tackles these challenges by allowing for replication of experimentally obtained values and trends. It thus helps to link the dynamical phenomena observed in vitro to the detailed atomistic motions, thereby assisting the interpretation of NMR results and offering greater insight into the biophysics of lipid membranes.

## AUTHOR CONTRIBUTIONS

M.D. and M.F.B. developed the concept and designed the research. G.K. produced the simulation trajectories and M.D. performed all computational analysis. M.D., G.K., and M.F.B. wrote and edited the manuscript.

### ACKNOWLEDGMENTS

M.D. is supported by NIH postdoctoral fellowship F32GM134704. G.K. is supported by the HRH Prince Alwaleed Bin Talal Bin Abdulaziz Alsaud Institute of Computational Biomedicine at Weill Cornell Medical College through the 1923 Fund. M.F.B. acknowledges NIH support through grant R01EY026041 and NSF through grants MCB 1817862 and CHE 1904125. Molecular dynamics simulations were performed using the Oak Ridge Leadership Computing Facility (Summit allocation BIP109) at the Oak Ridge National Laboratory (supported by the Office of Science of the U.S. Department of Energy under Contract No. DE-AC05-00OR22725).

## DECLARATION OF INTERESTS

The authors declare no competing interests.

## Notes

### Competing Interest Statement

The authors have declared no competing interest.

### Summary of Updates

(1) changing the title to more accurately capture the content (2) direct calculation of the angular dependence of the relaxation from the simulations, and the corresponding model-free orientationally averaged relaxation rates; (3) detailed theoretical description of the recently introduced approach published in 2023, and how it relates to the orientation-independent relaxation rate commonly calculated in MD studies; (4) new simulations with an output interval of 4 ps which allow for calculation of the orientation-dependent relaxation rates without any special fitting of the correlation functions; (5) robust comparison between the different types of relaxation rates, including their sensitivity to the resolution of the MD simulation data; (6) new analysis of the isomerization rates of the carbon segments on the lipid acyl chains; (7) expanded discussion on the orientation-independent relaxation rates and what prevents them from capturing the full range of dynamics of lipid CH bond dynamics in bilayers.

